# A CRISPR-Cas assisted shotgun mutagenesis method for evolutionary genome engineering

**DOI:** 10.1101/2021.09.08.459399

**Authors:** Ming Zhao, Miaomiao Gao, Liangbin Xiong, Yongjun Liu, Xinyi Tao, Bei Gao, Min Liu, Feng-Qing Wang, Dongzhi Wei

**Affiliations:** State key Lab of Bioreactor Engineering, Newworld Institute of Biotechnology, East China University of Science and Technology, Shanghai 200237, China; Shanghai Key Laboratory of Molecular Imaging, Shanghai University of Medicine and Health Sciences, Shanghai 201318, China

**Author notes:** Corresponding authors. Address: East China University of Science and Technology, P.O.B.311, 130 Meilong Road, Shanghai 200237, China., E-mail address (F. Wang); (D. Wei).

**Keywords:** CRISPR-mediated genome engineering, whole genome mutagenesis, iterative evolution

## Abstract

Genome mutagenesis drives the evolution of organisms. Here, we developed a <u>C</u>RISPR-Cas <u>a</u>ssisted <u>r</u>andom <u>m</u>utation (CARM) technology for whole genome mutagenesis. The method leverages an entirely random gRNA library and SpCas9-NG to randomly damage genomes in a controllable shotgun-like manner that then triggers diverse and abundant mutations via low-fidelity repair. As a proof-of-principle, CARM was applied to evolve the capacity of *Saccharomyces cerevisiae* BY4741 to produce β-carotene. After seven rounds of iterative evolution over two months, a β-carotene hyper-producing strain, C7-143, was isolated with a 10.5-fold increase in β-carotene production and 857 diverse genomic mutants that comprised indels, duplications, inversions, and chromosomal rearrangements. Transcriptomic analysis revealed that the expression of 2,541 genes of strain C7-143 were significantly altered, suggesting that the metabolic landscape of the strain was deeply reconstructed. In addition, CARM was applied to evolve the industrially relevant *Saccharomyces cerevisiae* CEN.PK2-1C, the S-adenosyl-L-methionine production of which was increased to 2.28 times after just one round. Thus, CARM is a user-friendly and practical strategy for genetic remodeling and reverse engineering to investigate complicated organismal metabolism.

## Introduction

Laboratory evolution strategies that emulate natural evolution are efficient mechanisms to rapidly promote the development of species with desired characteristics (Sandberg *et al*, 2019; Fernández-Cabezón *et al*, 2019; Yang *et al*, 2019). Gleizer *et al*. (Gleizer *et al*, 2019) and Gassler *et al*. (Gassler *et al*, 2020) recently successfully converted *Escherichia coli* and *Pichia pastoris* from heterotrophs to autotrophs using laboratory evolutionary methods. These significant advances have demonstrated that laboratory evolution is a powerful means to fully remodel the genetic and metabolic phenotypes of organisms that then provide unique opportunities to explore the metabolic potentials of living organisms, understand their complicated metabolic pathways, and develop powerful biological tools for industrial applications.

Genomic mutagenesis drives evolution. Under natural conditions, the spontaneous mutation rate in genomes is extremely low (typically 10^−10^–10^−9^ occurred at a site), which cannot support the temporal requirements for laboratory evolution needed to generate sufficient and rapid genomic variation (Fernández-Cabezón *et al*, 2019). Therefore, it is necessary to develop efficient mechanisms to promote genomic mutagenesis to successfully conduct laboratory evolution (Sandberg *et al*, 2019; Fernández-Cabezón *et al*, 2019; Yang *et al*, 2019; Jiang *et al*, 2020; Chen *et al*, 2020). Induced mutagenesis is an effective means to accelerate random mutations within genomes, which has been used for biological breeding (i.e., mutation breeding) since the 1920s. Since then, as a typical method of laboratory evolution, a variety of induced mutagenesis techniques have been subsequently developed (Fernández-Cabezón *et al*, 2019; Wang *et al*, 2019; Yang *et al*, 2019), which have provided a critical foundation for the establishment of the modern biological industry and have significantly promoted progress in industrial production of antibiotics, organic acids, vitamins, pharmaceuticals, enzymes, and steroids (Agrawal *et al*, 1999; Cai *et al*, 2018; Jiang *et al*, 2017; Zhang *et al*, 2018; Zhang *et al*, 2019; Zhu *et al*, 2018). Indeed, induced mutagenesis is still an important means for biological breeding in many enterprises and laboratories.

Nevertheless, the drawbacks of traditional induced mutagenesis methods are increasingly problematic (Fernández-Cabezón *et al*, 2019). Generally, the mutation types induced by a single mutagen often lack diversity, limiting their iterative application. Thus, it is often necessary to utilize multiple mutagens to achieve satisfactory mutation effects, rendering the process time and labor consuming. In addition, most mutagenic factors, especially chemical mutagens and radiation, are strongly toxic carcinogens. These intrinsic deficiencies have led to reduced use of mutagenesis, particularly with the development and widespread availability of genome editing technologies. Nevertheless, induced mutagenesis is still a highly practical means of laboratory evolution, especially under unknown genetic backgrounds, unclear metabolisms, or the lack of effective genome modification techniques. Consequently, the development of new genomic mutagenesis techniques like induced mutagenesis, but without its intrinsic deficiencies, is highly desirable.

Powerful alternatives to induced mutagenesis have gradually emerged since 2000, including genome replication engineering-assisted continuous evolution (GREACE) and multiplex automated genome engineering (MAGE) (Luan *et al*, 2013; Alper *et al*, 2006; Park *et al*, 2003; Wang *et al*, 2009). GREACE is an effective random mutagenesis method that can trigger automatic and continuous genomic mutagenesis by destroying replication fidelity. However, the poor stability of mutants that require complex and strict regulatory systems to control replication fidelity render extensive applications difficult (Fernández-Cabezón *et al*, 2019; Luan *et al*, 2020). MAGE is a precise genome mutation technique that uses synthetic oligonucleotides (30–110 bp) to introduce mutants into genomes via low-fidelity annealing with the lagging strand at a DNA replication fork. For organisms with highly active DNA mismatch repair activities, MAGE is not an effective genomic mutagenesis method, and its mutagenesis efficiency is closely related to the target distance from the origin of replication, wherein longer distances lead to less efficiency (Wang *et al*, 2009; Barbieri *et al*, 2017). The desirable performances of these methods for genome mutation suggests that developing genome engineering methods for inducing mutagenesis can be a potential mechanism to generate a large number of mutations in genomes that would enable laboratory evolution experiments.

The unprecedented success of CRISPR technology for gene editing in recent years has led to the development of multiple CRISPR-mediated genome engineering (CMGE) strategies like CREATE (Garst *et al*, 2017), CHAnGE (Bao *et al*, 2018), MAGESTIC (Roy *et al*, 2018), Istop (Billon *et al*, 2017), and EvolvR (Halperin *et al*, 2018). These novel CMGEs exhibit powerful abilities to alter genes at the genome-wide scale. Among these, CREATE, CHAnGE, and MAGESTIC are trackable and accurate genome editing methods that can massively introduce parallel mutants in genomes by array-synthesized gRNA coding sequences and homology-directed repair cassettes (Garst *et al*, 2017; Bao *et al*, 2018; Roy *et al*, 2018). Compared to traditional induced mutagenesis, CMGE generation attempts to achieve higher accuracy and traceability of mutagenesis with the aid of homologous recombination (HR) rather than genome variation complexity and diversity. iSTOP and EvolvR are two HR-independent genome engineering methods that do not require synthetic HR donor oligonucleotides for genome mutagenesis. Nonetheless, iSTOP only allows mutation-based conversion of four codons into STOP codons in the genome to facilitate gene inactivation (Billon *et al*, 2017), while the advantage of EvolvR is to diversify nucleotides in a user-defined window up to 350 nucleotides (Halperin *et al*, 2018). Although these existing CMGEs are distinct from induced mutagenesis methods, their high efficiency for genomic mutagenesis indicate that they may be viable platforms to develop a CMGE-like technology that could exhibit similar performance as induced mutagenesis for genome-wide scale mutagenesis, but with greater simplicity, flexibility, universality, and lower application costs.

Genomic damages caused by mutagens and low-fidelity repair are two key aspects of induced mutagenesis. In this study, we leveraged the basic mechanism of induced mutagenesis to develop a CRISPR-assisted random mutation (termed CARM) method that can simulate induced mutagenesis to introduce diverse mutations into genomes. To generate random genome-wide damage, a completely random guide RNA (gRNA) library was designed to guide Cas nuclease to randomly target genomes in a shotgun fashion. The DNA damages introduced by Cas nuclease is double-strand breaks (DSBs), a serious type of DNA damage. Consequently, mutations will be introduced at these DSB sites, in a manner similar to induced mutagenesis, by low-fidelity repair mechanisms for survival. As a proof-of-principle, CARM was applied in *Saccharomyces cerevisiae* to generate β-carotene producing strains. The method was then verified as an effective and user-friendly way to trigger diverse mutagenesis scattered around the entire genome, and thus could be readily used to promote iterative evolution in laboratory settings.

## Results

### Design of CARM for genome-wide shotgun mutagenesis

CARM was designed as a genome engineering technique based on a CRISPR/Cas system in order to generate shotgun-like mutagenesis and trigger random variation across the entire genome (Fig 1). To achieve this, an undifferentiated random mutagenesis approach at the genome-wide scale was required and accomplished via two means. First, a random gRNA library (gRNA-L) was constructed to cover all possible genomic loci and that could thus guide Cas nucleases towards random cleavage of targeted DNA across the entire genome. A protospacer adjacent motif (PAM) is necessary for Cas nucleases to recognize target loci and thus, as high as possible PAM density in the genome would clearly be necessary for a Cas nuclease to target as much as possible loci within a genome. Second, an engineered variant of *Streptococcus pyogenes* Cas9 (SpCas9-NG) with relaxed PAMs including NG, NAC, NTG, NTT, and NCG (Nishimasu *et al*, 2018), was employed. Using NG PAM as an example, its target frequency in a genome is equal to the GC content, and approximately one instance per 2 bp in a typical genome with around 50% GC. Considering other accessible PAMs besides NG and up to an 8 bp trimming window after cleavage at the −3 to −10 position relative to the PAM (Stephenson *et al*, 2018), SpCas9-NG, in combination with the gRNA-L as an editing system, was used to achieve an unprecedented random mutagenesis effect across the entire genome. To verify the feasibility of the above design, an engineered β-carotene-producing strain, C0 (Fig EV1), derived from *S. cerevisiae* BY4741 (Appendix Table S1) was used as the test model. Electroporation was used to randomly deliver hundreds of gRNA cassettes into a cell with the expression of SpCas9-NG, thereby guiding SpCas9-NG to randomly damage genomes in a high-throughput shotgun-like manner. The multiple DSB lesions generated in the genome would provoke complex cellular DNA repairs, while the cells without lethal mutagenesis would survive. Considering that CRISPR system creates DSBs within hours (Liu *et al*, 2020b), consequently, a widespread assault on the genome would rapidly be generated in a transformant. Then, the genome repair cells would grow into stable clones in the selected plate with deep metabolic remodelling, which can be screened by specific phenotypes for directed evolution, via, for example, the color of β-carotene in the study.

**Fig. 1.**
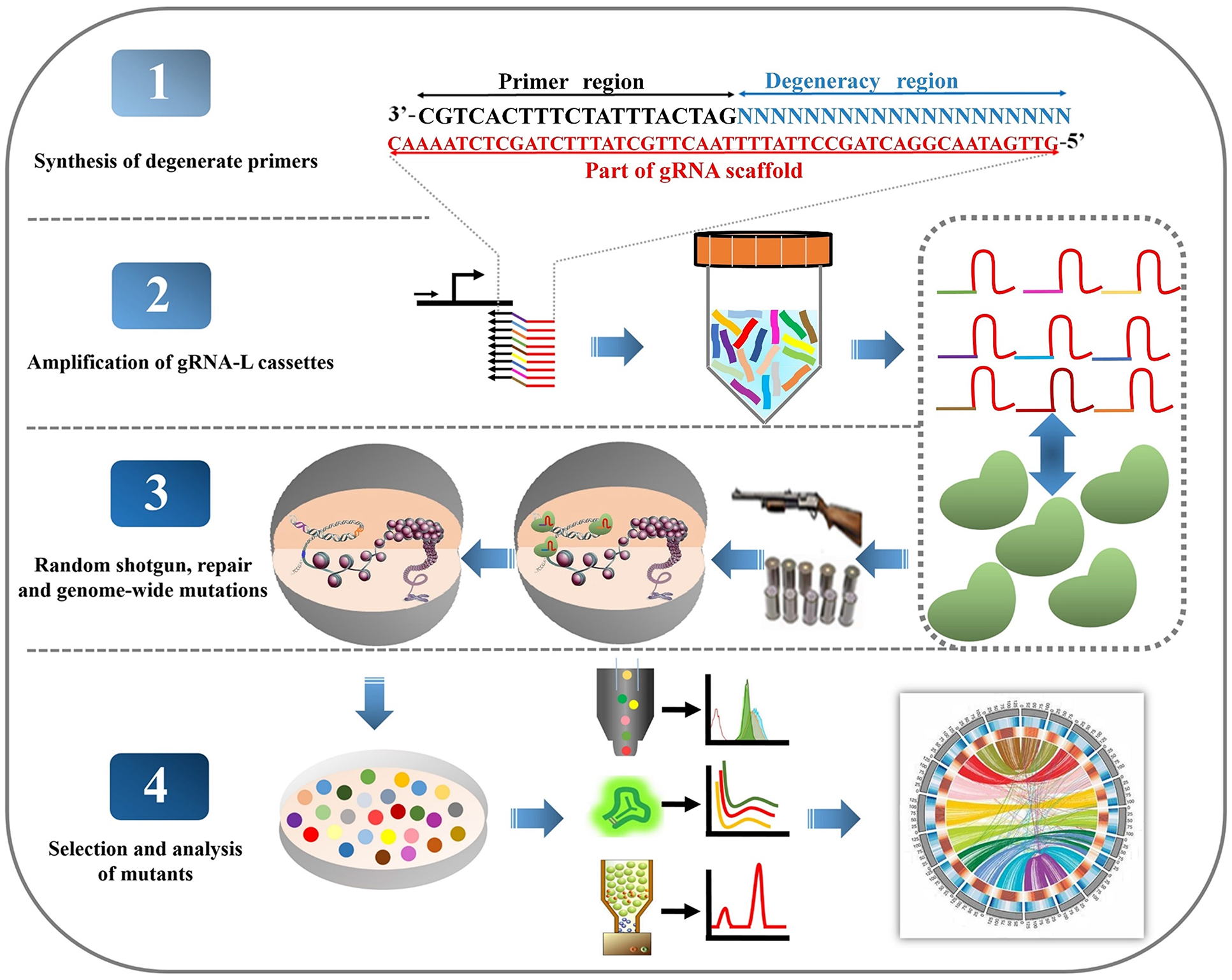
Design and schematic for CARM shotgun mutagenesis of genomes. A reverse primer library containing 20 degenerate bases was synthesized and used to amplify random gRNA library (gRNA-L) cassettes with a forward primer. The forward primer and the part of gRNA scaffold region contain over 50 bp homologous fragments with the ends of a linearized plasmid pSCM (Appendix Fig S2), respectively. After homologous recombination with the linearized plasmid pSCM, a number of different gRNA cassettes can be delivered into cells. By delivering random gRNA cassettes into a cell in conjunction with the expression of Cas proteins, random gRNAs are produced and instantaneously guide Cas proteins to break genomes in a random and high-throughput shotgun-like manner. During DNA emergency repairs, numerous and diverse mutations are triggered that can lead to deep metabolic remodeling of cells.

The construction of the gRNA-L is a critical priority in the design scheme (Fig 1). The genome size of strain C0 is approximately 12 Mbp. Thus, to cover all of the possible genomic loci, while not taking into account off-target effects and PAMs, the gRNA-L should contain at least 1.2×10^7^ gRNA. To synthesize a gRNA-L of this size would be time-consuming and labor-intensive. Consequently, we employed a simple and economical way to construct a greater gRNA-L by routine polymerase chain reaction (PCR) with primers that contain entirely degenerate guide sequences (20 bp) complementary to the target DNA. The amount of this kind of gRNA-L could reach 4^20^ (about 10^12^) in theory, which is much larger than the target genomes. Thus, the gRNA-L would be able to direct Cas nucleases to target almost any possible locus on a particular genome. Here, 4^20^ gRNAs is defined as a library unit (LU). In a typical PCR, 1 nmol (about 57 μg) of the degenerate primer, or about 6×10^14^ of the primer oligonucleotides, could be synthesized by the commercial phosphoramidite method and would only cost three US dollars. Approximately 3×10^12^ primer oligonucleotides were generally used to clone the expression cassettes of gRNA, and a gRNA-L with about 3 LU (∼1.5 μg) were readily achieved. To determine the randomization of the constructed gRNA-L, the amplified gRNA cassettes were sequenced using next-generation sequencing (NGS) and the 20 bp guide sequences of 198 gRNAs were statistically analyzed (Appendix Table S4). The guide sequences exhibited perfect degeneracy (Fig EV2), confirming that the method was an easy way to use degenerate PCRs to construct a sufficiently random gRNA-L to cover all possible loci within a genome.

Combining the gRNA-L and linearized gRNA expression plasmids led to the random delivery of the gRNA expression cassettes by electroporation into the SpCas9-NG-expressing strain C0 (Fig 2A and B). To generate a widespread assault on the genome, a high copy plasmid, pSCM, was used to carry a high number of gRNA cassettes. The number of gRNAs introduced into cells is a critical parameter that determines the assault effect of SpCas9-NG on a genome and the survival rate of cells. Consequently, we optimized and evaluated the transformation effects of gRNA cassettes into cells (Fig EV3). The results indicated that the number of surviving transformants containing the plasmids with gRNA cassettes were significantly lower than that without gRNA cassettes, demonstrating there were some lethal mutations. Considering that the higher mortality means the more damage occurred on the genome, the condition with the highest mortality up to about 35% (marked with red arrow in Fig EV3 was employed in this study. Under the selected conditions (Appendix Table S5), about 130 gRNAs could be delivered into a single cell on average (Fig EV4). Thus, numerous DSBs could be introduced into the whole genome, leading to low-fidelity repairs that would result in diverse mutation loci.

**Fig. 2.**
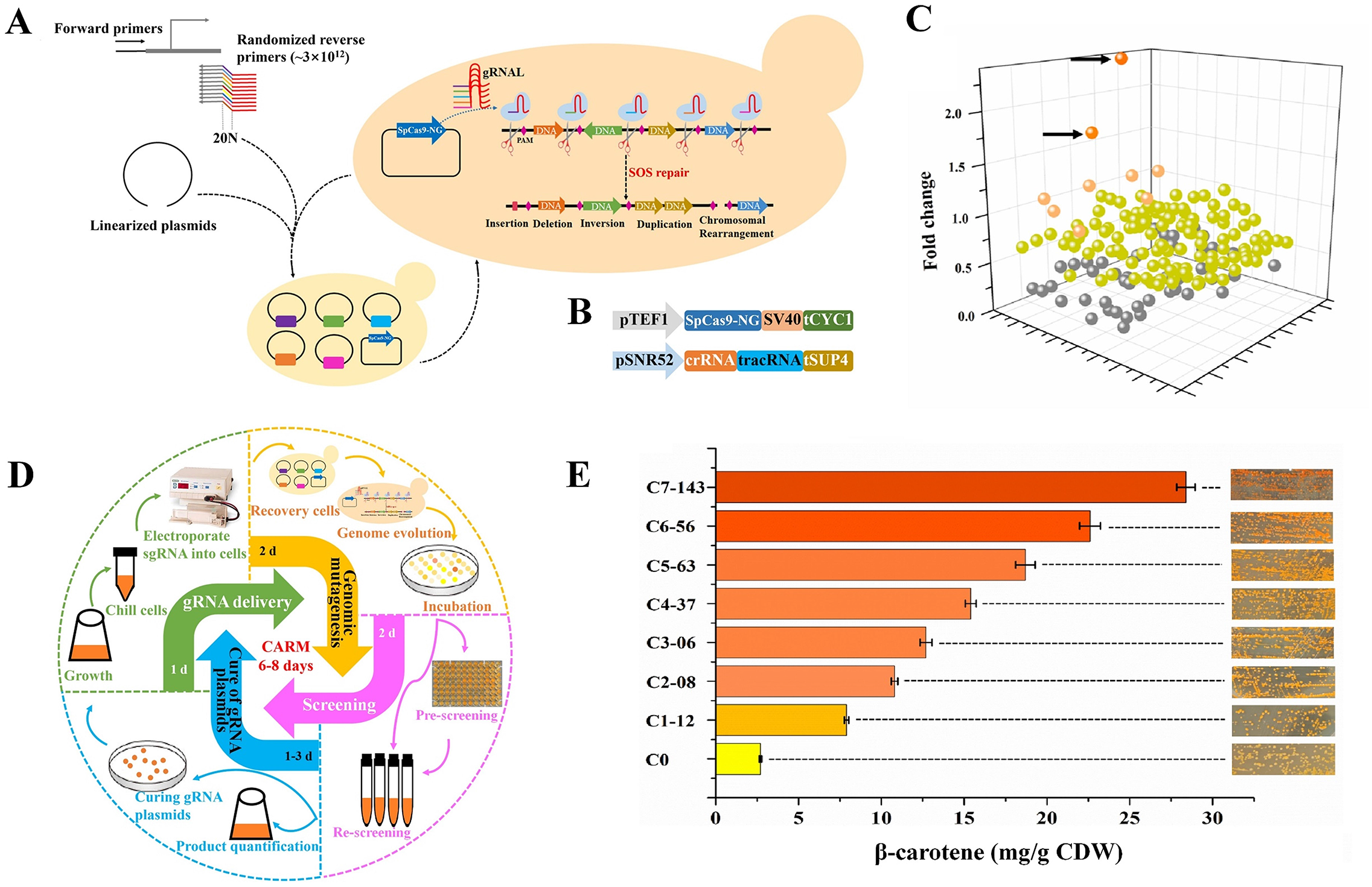
Construction and evaluation of CARM for driving rapid evolution of *S. cerevisiae*. A CARM construction in *S. cerevisiae*. B Regulation of CARM in *S. cerevisiae*. C Positive mutation rate analysis of *S. cerevisiae* C0 treated by CARM for β-carotene production. D Schematic of the iteration CARM cycle. E Analysis of β-carotene production of screened mutants by iterative evolution. Error bars show standard deviation from three independent experiments.

### Effectiveness of CARM for generating cellular variation and evolution

DSBs are the most dangerous type of genomic damage and their repair is essential for cellular survival. Without exogenous homologous recombination donors, multiple error-prone DNA repairs play key roles that introduce diverse non-lethal mutants at the DSBs in resurgent cells (Lian *et al*, 2019; Maslowska *et al*, 2018). To determine if CARM can effectively trigger diversified genomic variation, as guided by hundreds of gRNAs, a laboratory evolution experiment of *S. cerevisiae* strain C0 to increase the production of β-carotene was performed (Fig 2A and B).

The effects of CARM on the production of β-carotene was first evaluated. 176 viable colonies were indiscriminately isolated from C0 treated by CARM and their β-carotene yields were determined without regard to colony colors on screening plates. Among these strains, two mutants exhibited a >20% increase in β-carotene production in comparison with strain C0, and fifty mutants exhibited a >50% decrease (Fig 2C). Strain C0 transformed with gRNA-unloaded pSCM was used as a control and exhibited no changes in the production of β-carotene (Fig EV5). The changes in β-carotene production clearly demonstrated that CARM altered the metabolic status of cells. Different β-carotene contents in cells can cause colonies to exhibit different orange colorations and so colonies with deeper β-carotene colors than strain C0 were enumerated on ten screening plates. The number of positive variants was around 1% of the viable strains. A total of 20 colonies (from about 10^4^ colonies) were subsequently picked based on deep β-carotene coloration. The selected strains’ yields of β-carotene were further verified by submerged fermentation in shake flasks. Among the strains, strain C1-12 was selected due to its strong capacity for β-carotene production (7.9 mg/g cell dry weight (CDW)), which was 2.92 times that of strain C0 (Fig 2E, Fig EV6 and EV7). Thus, these results demonstrate that CARM can be used as an effective mechanism to promote laboratory evolution.

To further investigate the applicability of CARM for laboratory evolution, iterative CARM was conducted to further boost the production of β-carotene (Fig 2D). The process comprised four steps: gRNA delivery and genomic mutagenesis, preliminary screening, secondary screening, and curation of gRNA plasmids. Six rounds of iterative evolution were then performed using strain C1-12, resulting in strains C2-08, C3-06, C4-37, C5-63, C6-56, and C7-143 that were screened in each iterative cycle based on >20% increases in β-carotene production (Fig 2E, Fig EV6 and EV7). The β-carotene production of the final strain, C7-143, reached 10.5 times that of strain C0, with yields reaching up to 28.4 mg/g CDW. The six iterative cycles of CARM were achieved within two months and a complete CARM cycle took about 6–8 days, on average. Therefore, CARM held promise to be effective, user-friendly, and rapid for inducing laboratory evolution, and they could circumvent shortcomings of induced mutagenesis and CMGE.

### CARM profile for genome-wide mutagenesis and metabolic remodeling

CARM was designed to trigger random and diverse mutagenesis activities at the genome-wide scale via random gRNA library-guided SpCas9-NG. The above results demonstrate that the use of the CARM method can lead to the delivery of hundreds of gRNA into *S. cerevisiae* and thus promote its phenotypic evolution towards increased β-carotene production, implying that the expected genomic variation was triggered. To confirm the genomic evolution, the genomes and transcriptomes of strains C3-06, C5-63, C7-143, and C0 were sequenced and investigated. The primary genomic variation introduced by nonhomologous DNA repairs at DSB sites generally comprise insertions or deletions of short DNA fragments, wherein the repairs of adjacent DSBs likely trigger the deletion, inversion, or rearrangement of larger fragments. All of the observed variation types, with the exception of single base pair substitutions, were statistically evaluated using structural variation analysis. Compared to the C0 strain, 422, 681, and 857 mutations were detected in strains C3-06, C5-63, and C7-143, respectively (Fig 3A). On average, each cycle of CARM generated about 122 variants during the evolution of β-carotene producing strains. Mapping these variations to the C0 genome indicated that mutation numbers increased one after another in the genomes of C3-06, C5-63, and C7-143 and were distributed across all of the 16 yeast chromosomes without any particular regularity, demonstrating that the mutagenesis is random. Among these mutations, indels of short fragments (< 10 bp) dominated, comprising more than 84% of the total mutations. In addition, some indels of large fragments and chromosomal rearrangement events were also identified among strains. For example, 27 indels of >100 bp, including two duplications and one inversion, and 13 chromosomal rearrangements were present in strain C7-143 (Fig 3B and C, Appendix Table S6). These data confirmed that the genome of *S. cerevisiae* accumulated a large number of mutations via DNA cleavage and error-prone repairs under the guidance of hundreds of gRNAs. Thus, these data also confirmed that CARM is an effective measure to trigger random and diverse mutagenesis across the entire genome in a shotgun-like manner.

**Fig. 3.**
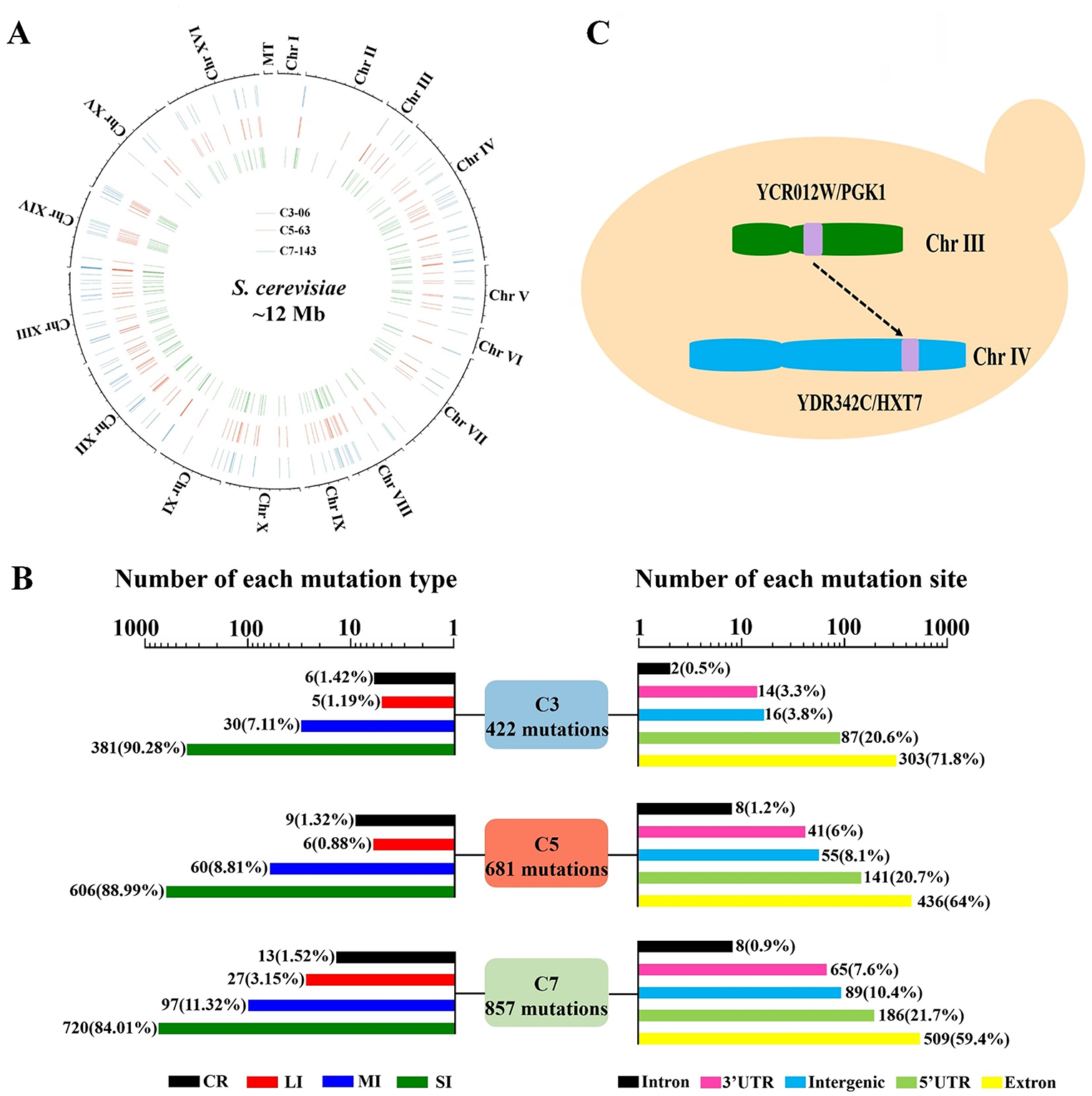
Genomic analysis of evolved strains. A Circos map showing the genetic changes of evolved strains relative to the parent strain. The blue, red, and green bars show genetic changes of evolved strains C3-06, C5-63, and C7-143, respectively. B Number and types analysis of mutations. CR represents chromosomal rearrangements; LI represents long fragment (>100 bp) DNA indels; MI represents medium fragment (10-100 bp) DNA indels; SI represents short fragment (<10 bp) DNA indels. C A sample of CR occurred in C7-143.

Interestingly, the expression cassettes of the exogenous genes *crtE*, *crtYB*, and *crtI* introduced into the genome of C0 to biosynthesize β-carotene did not exhibit any mutations. Thus, the significant increases of β-carotene production by these mutants can be attributed to profound changes of the basic host metabolism instead of through the introduced metabolic pathway. Further analyses indicated that about 40% of the mutations in strain C7-143 occurred in the non-coding (Fig. 3B), though the abundance of this region is 27.2% of the yeast genome (Alexander *et al*, 2010). Apart from that, the mutation frequencies in the other locus types did not exhibit significant differences. The enrichment of these mutations in the untranslated regions might be attributed to the possibly higher lethal effects of mutagenesis at other loci, such as the exons of essential genes. In addition, the enrichment of these mutations in the untranslated regions and the decrease of these mutations in the intergenic region might be attributed to the directed screening of β-carotene producing strain. Furthermore, the results also signified that the host metabolism likely underwent intensive changes. To confirm this hypothesis, transcriptomic profiles in strains C3-06, C5-63, and C7-143 were evaluated in comparison with those of the original strains C0 and BY47419_NG_ at the exponential growth metaphase with the same growth status (18 h). Compared to strain BY47419_NG_, the engineered strain C0 harbouring an exogenous β-carotene biosynthesis pathway exhibited minimal changes in global transcription profiles, with just 55 upregulated genes and 42 downregulated genes, among about 5,893 total genes (Fig 4A and B). Thus, the introduction of the exogenous genes for β-carotene biosynthesis did not exert a high impact on the global metabolism of the host. In stark contrast, the transcriptional patterns of strains C3-06, C5-63, and C7-143 exhibited considerable differences to those of strain C0, with 1,148, 1,285, and 1,509 upregulated genes and 889, 1,072, and 1,290 downregulated genes (|Log2FoldChange|>1; *p* < 0.05, Fig EV8), respectively. These data clearly demonstrate that the global metabolism of the host underwent tremendous changes after CARM-based evolution. Further, these observations highlight the significant value of CARM for deep reconstruction of cellular metabolism by iterative mutation and evolution.

**Fig. 4.**
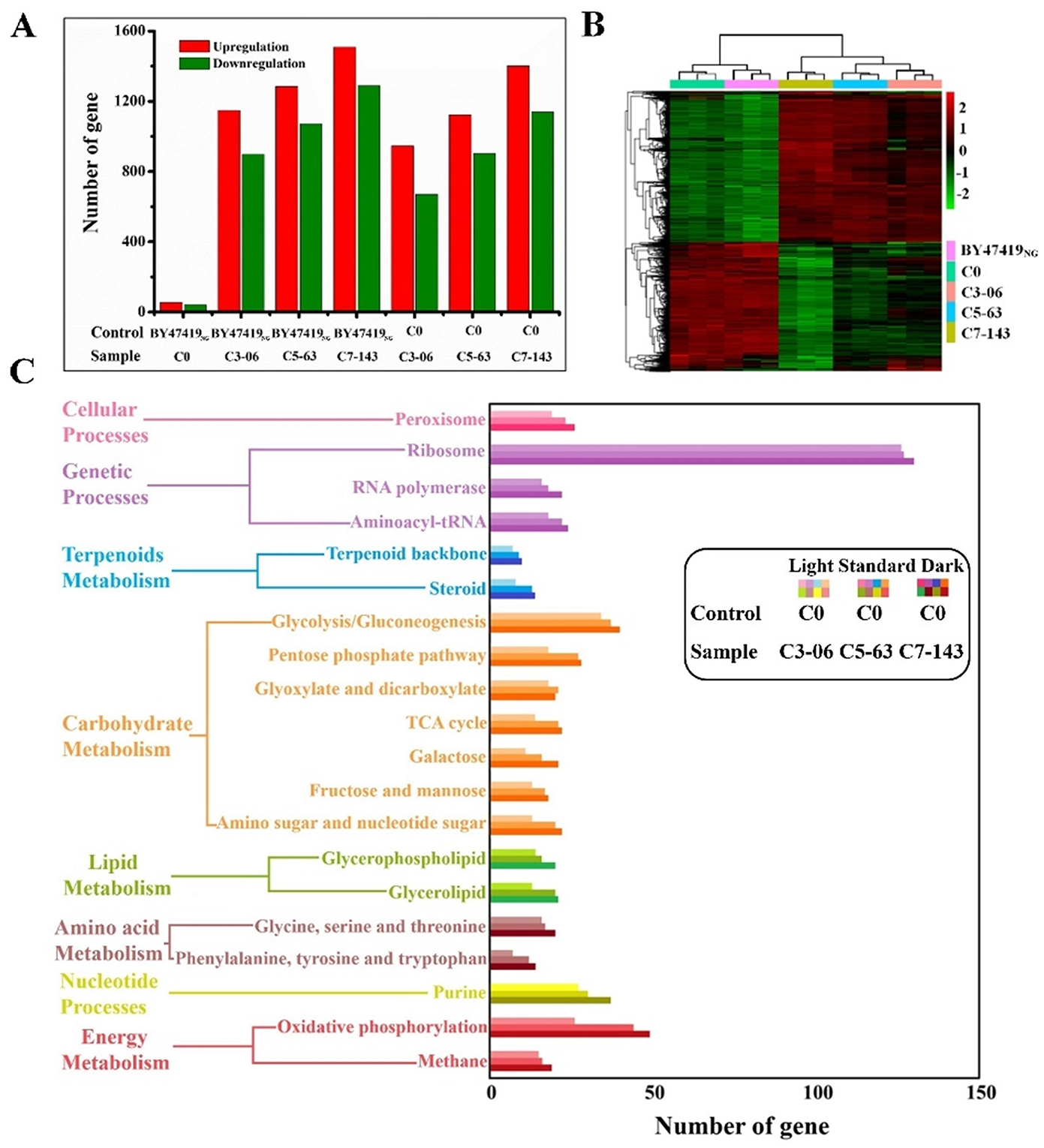
Transcriptomic profiling of evolved strains. A The number of genes significantly (|Log2FoldChange|>1; *p* < 0.05) differentially expressed in the evolved strains C3-06, C5-63, and C7-143, relative to control strains. B Heatmap of transcript diversity across evolved strains. C Functional enrichment analysis of transcripts based on KEGG pathways. All strains were grown in YPD media at 30°C until logarithmic phase. A–C Three biological replicates were performed per strain.

The biosynthetic capacity of β-carotene was used as a screening marker and thus, the changes in metabolism of these mutant strains should be conducive to producing β-carotene. To investigate this hypothesis, KEGG (Kyoto encyclopedia of genes and genomes) pathway enrichment analysis was used to identify metabolic differences between the evolved strains and the original strain, C0. Changes in ribosomal pathways stood out among the metabolic variations. Among a total of 158 genes involved in ribosomal pathways, more than 125 genes exhibited upregulated transcriptional levels in strains C3-06, C5-63, and C7-143 (Fig 4C), suggesting that deep changes occurred in the global gene translation networks of these evolved strains. Thus, such mechanisms may be pivotal factors in the global change of β-carotene biosynthesis metabolic phenotypes. It is worth noting that significant changes also took place in the glycolysis and TCA cycle pathways. Interestingly, some key genes of the glycolysis pathway were upregulated, while some key genes of the TCA cycle were significantly downregulated (Appendix Table S7), indicating that additional acetyl-CoA precursor could be available for β-carotene biosynthesis in these evolved strains. In addition, key genes for terpenoid backbone biosynthesis were also significantly upregulated after evolutionary screening. Thus, these results point to the evolution of a specific pathway to increase the biosynthesis of β-carotene. In this pathway, acetyl-CoA supply from a carbon source, such as glucose, is increased by increasing glycolysis activity, but weakening TCA cycle activity allows the conversion of more acetyl-CoA to β-carotene via the enhancement of the terpenoid biosynthesis pathway. In addition, marked gene expression differences were also noted for lipid metabolism, and especially in the biosynthesis of glycerophospholipid (14/39, differentially expressed genes number / total genes number) and ergosterol (13/17) (Appendix Table S7), both of which are essential components for cellular membranes. The expression of genes related to peroxisome synthesis were also enhanced in these evolved strains, consistent with our previous research that peroxisomes being a storage space for hydrophobic terpenes that are necessary for β-carotene synthesis in *S. cerevisiae* (Liu *et al*, 2020a; Li *et al*, 2020). We further confirmed that β-carotene accumulated in peroxisomes and that peroxisome numbers significantly increased with the trajectory of evolutionary selection (Fig EV9).

The above genomic variation and transcriptomic profiling data thus confirmed that CARM is a highly efficient way to trigger deep changes in cellular metabolic phenotypes by random and diverse genomic mutagenesis. Importantly, these mutagenesis effects can effectively be used to conduct laboratory evolution experiments or reverse engineer strains to investigate sophisticated relationships between genetic and metabolic phenotypes.

### Production performance and stability of evolved strains

An important potential application of CARM is as an alternative to mutation breeding to develop mutant strains with desired performances for industrial production or genotype-phenotype mapping. Genetic instability is a common concern for induced mutants and thus, the stability of strains developed with CARM was further examined using C7-143 as an example. After curing the plasmids carrying SpCas9-NG and gRNA, C7-143 was successively propagated for 10 generations and their colony morphologies and β-carotene production variation were investigated (Fig EV10). Changes in colony morphology and decreased β-carotene production were not observed, suggesting the presence of stable inheritable characteristics in *Saccharomyces cerevisiae* produced by genome mutagenesis via CARM.

The fermentation ability of C7-143 was then evaluated in a 3 L fermentator through fed-batch way, as previously reported (Xie *et al*, 2014). β-carotene production was closely related to cell growth, with the final titer of β-carotene reaching 1.1 g/L in four days, or about 26.1 mg/g CDW (Fig 5A and B). This value represents one of the highest outputs of β-carotene achieved in *Saccharomyces cerevisiae* without traditional metabolic pathway optimization (Xie *et al*, 2014; Cheng *et al*, 2020; Bu *et al*, 2017; López *et al*, 2019). In addition, to verify the generality of CARM, CARM was applied for the strain improvement to enhance the production of S-adenosyl-L-methionine in the industrially relevant *Saccharomyces cerevisiae* CEN.PK2-1C (Fig 5C). Combined with the efficient screening strategy for the SAM high-yield strain, the evolutionary strains, engineered after first round of CARM, were plated on the SD-Leu-Ura medium containing 0.4 mmol/L ethionine, and the largest colony, C1-SAM, was screened and cultivated in the optimized medium in the shake flask. The result showed that the SAM yield of evolutionary strain C1-SAM, reaching about 59 mg/L, was increased to 2.28 times that of wild strain CEN.PK2-1C (Fig 5D). Moreover, C1-SAM exhibited the faster growth rate than wild strain in the YPD medium containing 0.4 mmol/L ethionine (Fig 5E), which indicated that C1-SAM could produce much more SAM to resist the toxicity of ethionine. In summary, the performance of strain C7-143 and C1-SAM further demonstrated that CARM is a simple and efficient method to conduct laboratory evolution and rapidly develop robust strains for potential industrial application.

**Fig. 5.**
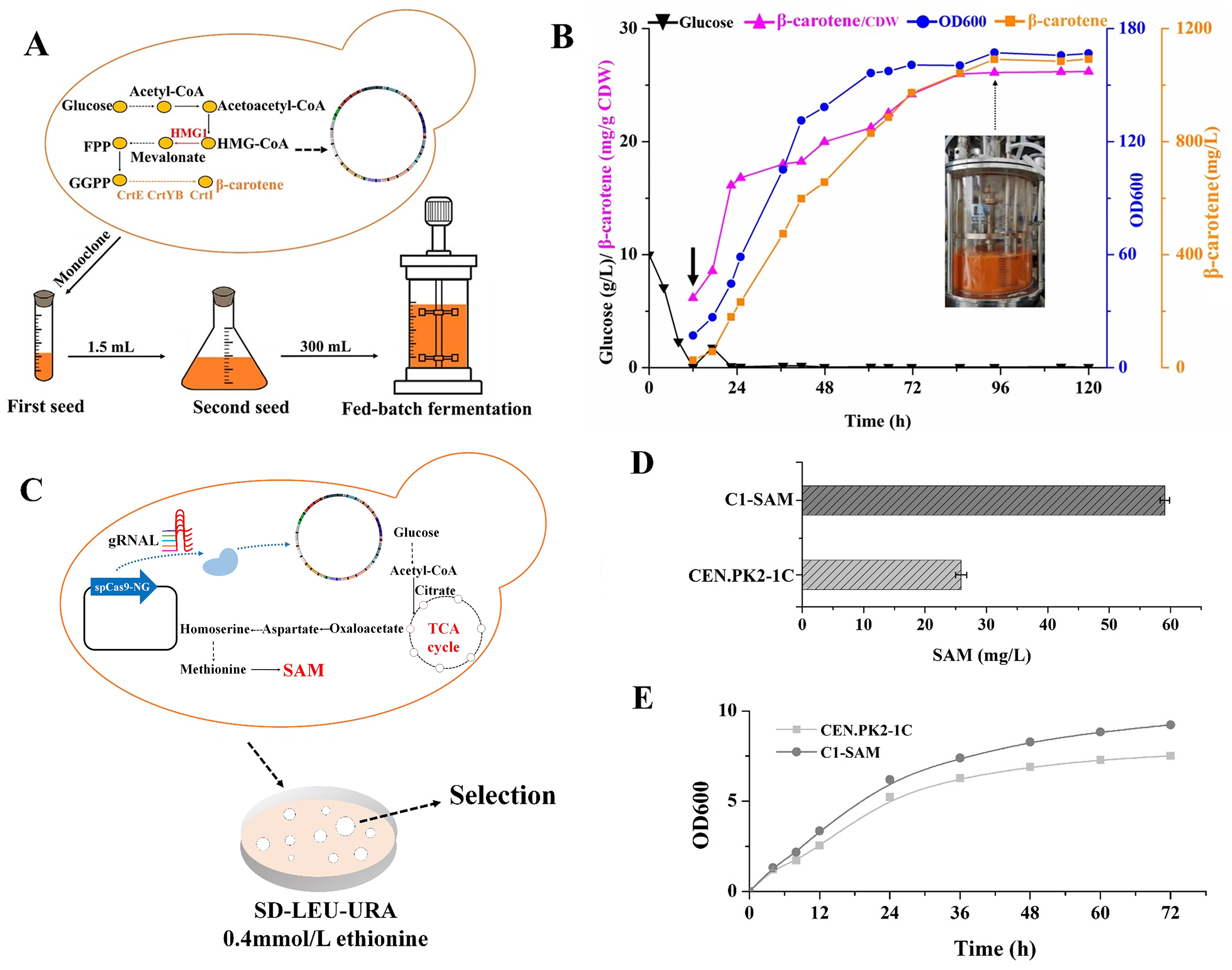
High-density fermentation of C7-143 in a 3 L batch bioreactor to stimulate β- carotene production. A Schematic diagram of the process of fed-batch fermentation. B High-density fermentation for β-carotene production. Time courses show changes in cell growth and β-carotene production of strain C7-143. The black arrow shows the point at which glucose was added to the medium. C Schematic diagram of the evolution and selection of SAM higher production strain. D Analysis of SAM production of screened mutant. E Growth curves analysis of screened SAM higher production strain in the YPD medium containing 0.4 mmol/L ethionine D and E Error bars show standard deviation from three independent experiments.

## Discussion

Compared to natural evolution, induced mutagenesis via physical and chemical means exhibits extensive potential to generate genome variation and evolution in laboratory settings. However, traditional mutagenesis methods have become more and more obsolete, due to the limited mutation diversity, the low mutation probability at the genome-wide scale, hazards to people and environments, and the requirement for special equipment, mutagens, or conditions. In contrast, genome engineering strategies like CMGE methods exhibit high efficiency and safety, but are also limited by some factors including high costs, the inability to generate diverse random mutations, and insufficient genomic coverage of mutations. Herein, we developed a novel induced mutagenesis method, CARM, that is based on CMGE technology, but can induce mutations in a high-throughput manner. Specifically, CARM combines the advantages of both traditional induced mutagenesis and CMGE to generate genomic variation, while exhibiting greater simplicity, safety, cost efficiency, mutation rates, and mutation type diversity to drive genomic mutagenesis. Consequently, CARM can be used as a powerful tool to perform iterative laboratory evolution experiments or to map complex genotype-phenotype relationships.

Two unique approaches were adopted in CARM to achieve multiplex random mutagenesis at the genome-wide scale. First is the use of the CRISPR-associated protein, as a tool to damage DNA, producing similar effects as mutagenesis via traditional physical and chemical mutagens. To achieve highly random mutagenesis across the whole genome, SpCas9-NG with the most minimal NG PAM requirements and other noncanonical PAMs, including NAC, NTT, and NCG, were used to achieve a high density of PAM targets distributed throughout the entire genome. Recently, a PAMless CRISPR-Cas9 variant SpRY has been engineered, which can target any loci of the genome nearly without PAM constraint (Walton *et al*, 2020). So, it is conceivable that the employment of SpRY or other PAMless Cas proteins will make CARM more feasible to induce random mutagenesis in a genome-wide scale. Second is the utilization of a gRNA library, guiding SpCas9-NG to randomly assault genomes at numerous possible loci. The random gRNA library (referred to as gRNA-L) was constructed using conventional PCR with degenerate primers to generate completely random guide sequences of gRNA (Fig 1 and Fig 2A). The 1 LU capacity of gRNA-L reached 4^20^ (about 10^12^) gRNAs, which is considerably larger than the size of any genome, including *S. cerevisiae* (about 12 Mbp) used here. To cover all of the genomic sites of *S. cerevisiae*, only 1.2×10^7^ gRNAs are needed at most if no off-target effects are taken into account. Thus, about 10^5^ gRNAs in the gRNA-L contain only 1 gRNA that can precisely target the genome of *S. cerevisiae*. Surprisingly, over 100 *S. cerevisiae* genome loci were mutated at a time by CARM. However, individual cells could only accommodate on the order of about 130 gRNAs at a time. This clearly suggested that most of the mutations were actually triggered by off-target effects. CRISPR-Cas9 has been reported to exhibit a high off-target rate of up to 66% and that the off-target mutations are often observed at frequencies greater than the intended mutations (Fu *et al*, 2013; Zhang *et al*, 2015). Such high rate of off-target damage is due to multiple PAM-distal mismatches between guide RNAs and target DNA (Hsu *et al*, 2013). Recently, Newton et al. has disclosed that intracellular DNA stretching and bubbles can induce CRISPR/Cas9 off-target effects at cryptic sites containing up to ten mismatches, indicating that CRISPR/Cas9 is a more promiscuous genome editing system than previously thought (Newton *et al*, 2019). If considering up to ten mismatches, the capacity of the gRNA library actually needs to be just 4^10^ (1.68×10^5^) in order to meet the requirements for random mutagenesis by CARM. Accordingly, a gRNA from the gRNA library may target up to about 71 (1.2×10^7^/1.68×10^5^) loci in the yeast genome based on the 12 Mbp (1.2×10^7^ bp) genome of *S. cerevisiae*. Actually, the average mutation probability triggered by CARM was approximately 0.93 (122 mutations / 131 gRNAs), which is within the possible range of off-target effects. Indeed, off-target mutagenesis is a major concern for CRISPR-Cas9 use in precise genome editing in biomedical and clinical applications, but in the context of this study, it is a favorable feature that can be exploited for CARM to trigger abundant and diverse unexpected mutants in a genomic background.

Using the above design schemes, abundant genomic damage can be triggered by CARM through both on-target and off-target mutagenesis in a shotgun-like manner, thereby leading to the simultaneous introduction of high-throughput mutations into a genome. Obviously, the number of mutations depend on the number of gRNAs delivered into cell. Therefore, the mutation frequency can be tuned according the requirements by optimizing the transformation amount of gRNAs into individual cells. Consequently, in combination with a high-throughput screening method, CARM can be used as an effective measure for directed evolution of genomes. In this study, CARM was tested by introducing and evolving β- carotene production capacity of *S. cerevisiae* strain C0. After seven rounds of iterative evolution over two months, the β-carotene titer was rapidly increased to 10.5 times that of the origin strain C0 (Fig 2E). Compared with reported physical and chemical mutagenesis methods (Bhosale *et al*, 2001; Mehta *et al*, 2003), CARM system is much more efficient for increasing the β-carotene yield of evolutionary strains. Thus, these results demonstrated that CARM was indeed an effective, fast, economical, and practical method to drive genomic variation.

The successful application of CARM in *S. cerevisiae* warrants the pursuit of its extensive application in other organisms. Based on the proof-of-principle CARM application in *S. cerevisiae* described here, three key areas should be considered in its application in other organisms. First is the toxicity of SpCas9-NG in some cells. Thus, an appropriate Cas protein should be selected for the study organism. Second is the simultaneous transformation of a number of gRNAs into a cell at one time that can trigger high-throughput assaults of Cas proteins on genomes. Electroporation was used in this study to achieve these effects, although this may not be suitable for all cells, and another appropriate method should be employed, when needed. Third, the mechanism and propensity of mutation may need to be carefully considered in future studies. In this study, at least 857 mutations occurred in the genome of *S. cerevisiae* C0 after seven rounds of iterative evolutionary screening, including 84.0% of mutations being indels of < 10 bp, 11.3% as indels of 10 bp to 100 bp, 3.2% as indels of >100 bp, and 1.5% as chromosomal rearrangements (Fig 3B). Notably, single base pair substitutions were not included as possible mutations in this study, because the genome sequencing accuracy of third generation DNA sequencing (about 99.999%) may not be high enough to assure complete confidence in single base pair mutations. Thus, the genomic variations analyzed here were confidently assigned to the activities of low-fidelity DNA repair results of abundant genome DSBs triggered by CARM, when precluding homologous recombination donor templates. Non-homologous end joining (NHEJ), microhomology-mediated end-joining (MMEJ), and single strand annealing (SSA) are ubiquitous error-prone repair pathways of DSBs among different organisms that can generate diverse low-fidelity DNA repair results including insertions, deletions, duplications, chromosomal rearrangements, and other potential mutation events (Pathak *et al*, 2019; Yeh *et al*, 2019; Chang *et al*, 2017; Shou *et al*, 2018; Allen *et al*, 2019; Chen *et al*, 2019; So & Martin, 2019; Her & Bunting, 2018; Taheri-Ghahfarokhi *et al*, 2018). *S. cerevisiae* contains all three repair pathways (Jiang *et al*, 2020; Lee & Lee, 2007). Individual contributions of these repair pathways to the mutations induced by CARM in this study are unclear, and warrant further investigation. It is worth noting that not all organisms contain all of these three DNA repair pathways, with one example being the absence of the NHEJ pathway in *E. coli* (Chayot *et al*, 2010). Consequently, the application of CARM in other organisms may differ from that shown here for *S. cerevisiae*.

The mutagenesis mechanism of CARM is very similar in principle to traditional physical and chemical mutagenesis. The latter employ physical or chemical mutagens to introduce damage into genomes and then generates mutations after low-fidelity DNA repair. For instance, γ-ray mutation can enable DSBs or single-stranded breaks in genomes, thereby inducing diverse genomic variation comprising indels, translocations, inversions, DNA fragmentation, and chromosomal aberrations (Lotfy *et al*, 2007). CARM similarly results in such effects. The critical difference among these methods is that a CRISPR-Cas system is used in CARM to introduce random damage into the genome and induce genomic variation based on low-fidelity DNA repair. In contrast, lesion mutagenesis that is induced by physical or chemical mutagens is usually sequence-specific rather than random. For example, mutagenesis induced by ultraviolet radiation is primarily caused by the dimerization of adjacent pyrimidines in a genome. Likewise, mutagenesis triggered by alkylating agents primarily affect base transitions, such as transitions of G to C or A to T. Therefore, a saturation effect of mutagenesis will occur when using the same chemical or physical mutagenesis methods several times. Thus, to achieve desired mutagenesis effects, several mutagenesis methods are often needed in combination. In contrast, genomic mutagenesis induced by CARM is highly random due to its shotgun-like targeting of the genome, as guided by the combination of a random gRNA library and nearly PAMless Cas proteins (Nishimasu *et al*, 2018; Walton *et al*, 2020). Consequently, the saturation effect will be minimal when using CARM, and it can be successfully used in successive iterations.

CARM is a CMGE technology that significantly differs from existing CMGE technologies including CREATE, CHAnGE, and MAGSTIC (Garst *et al*, 2017; Bao *et al*, 2018; Roy et al, 2018; Billon *et al*, 2017; Halperin *et al*, 2018). First, CARM occurs independently of HR to introduce variation in a genome. Second, CARM employs a completely random gRNA library to guide Cas proteins, which would be universal for all organisms and can readily be constructed via routine PCR. In contrast, CREATE (Garst *et al*, 2017), CHAnGE (Bao *et al*, 2018), and MAGSTIC (Roy *et al*, 2018) depend on HR and synthesized guide-donor oligos that are designed by computer tools on the basis of precise genome sequences. CARM is thus a simpler and more inexpensive means to engineer genomes. CREATE (Garst *et al*, 2017), CHAnGE (Bao *et al*, 2018), and MAGSTIC (Roy *et al*, 2018) exhibit powerful application in producing trackable and accurate genome editing and excel in the mapping of mutations at a single-nucleotide resolution for protein, metabolic, and genome engineering. Conversely, CARM is a highly random and high-throughput genome mutagenesis technology that has the potential to excel in driving complex genome evolution.

Nevertheless, current CARM also has some shortcomings to be improved in the follow-up study. First, a large-scale plasmid library needs to be constructed despite that it was simplified in this study via the HR recombination of linearized plasmids with the gRNA cassette library in yeast cell. Second, the delivery way of the gRNA library needs to be changed accordingly in different hosts due to their distinct differences in genetic manipulation systems. Third, the congruent relationships between the mutation types and the non-fidelity repair pathways triggered by CARM are unclear, which deserve further investigation.

## Materials and Methods

### Reagents and Tool table

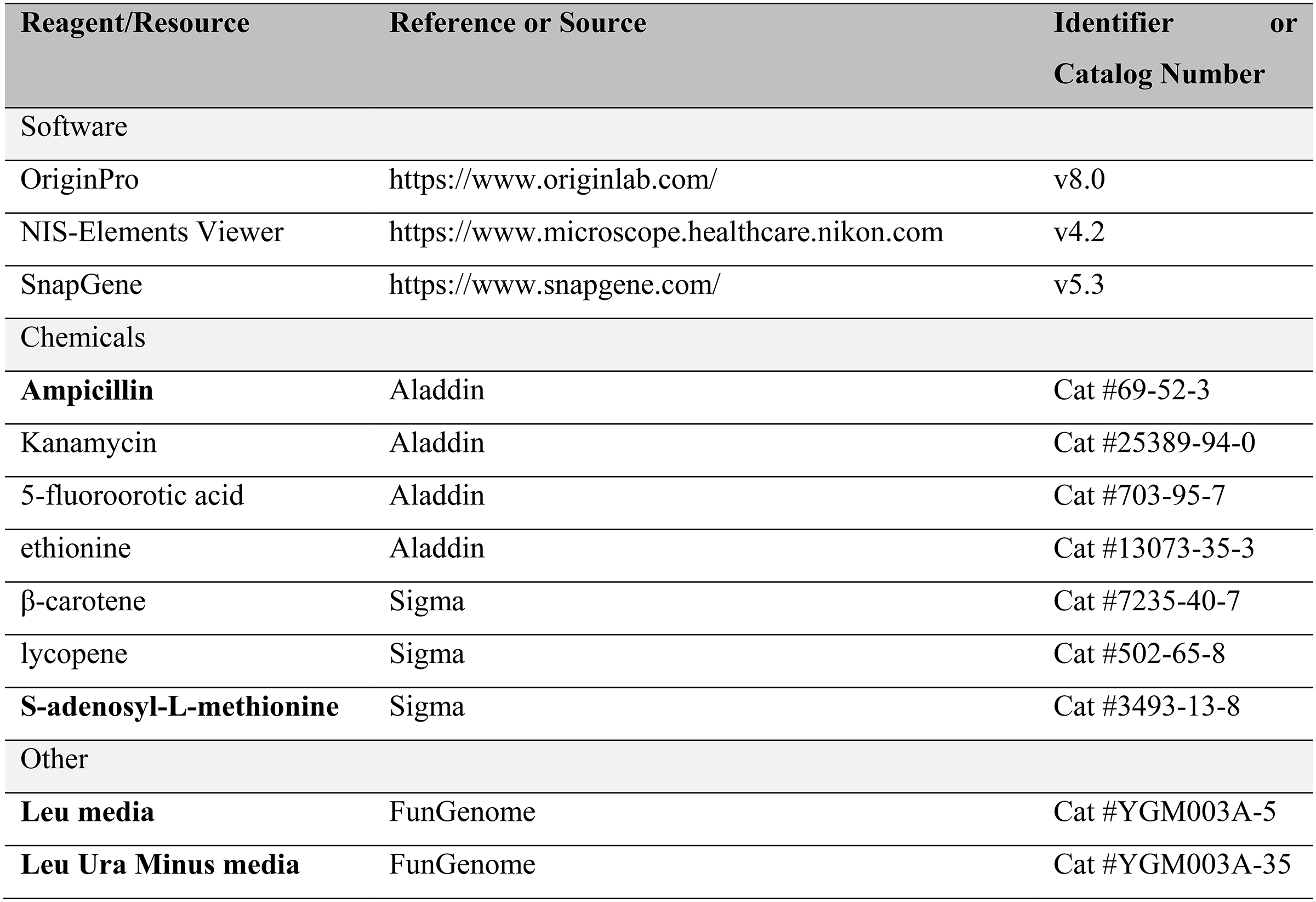

## Methods and protocol

### Strains, media and growth conditions

Strains used in this study are described in Appendix Table S1. *E. coli* DH5ɑ was used as the host for cloning and was grown aerobically on a rotary shaker (200 rpm) at 37°C in Luria-Bertani (LB) broth or on a plate containing LB supplemented with 1.5% (w/v) agar. *S. cerevisiae* strain BY4741 was used as the original host. In our previous study (Li *et al*, 2018), the key genes for the β-carotene biosynthetic pathway were introduced into the BY4741 genome. Here, the plasmids in the β-carotene producing strain were cured and then pCas9-NG was introduced into the strain to generate a C0 strain. Yeast strains were cultured in YPD (yeast peptone dextrose) broth medium (10 g/L yeast extract, 20 g/L peptone, and 20 g/L glucose) at 30°C. Synthetic complete drop-out medium with 2% D-glucose and without leucine (SD-Leu) and synthetic complete drop-out medium with 2% D-glucose and without leucine and uracil (SD-Leu-Ura) were used for cultivation and selection of yeast transformants. SD-Leu medium with 1 mg/mL 5-fluoroorotic acid (SD-Leu-5-FoA) was used for counterselection of guide RNA (gRNA) expression plasmids with the URA3 marker.

### CARM construction and the delivery of gRNAs into cells

Plasmids and primers that were used in CARM construction are shown in Appendix Table S2 and S3, respectively. pTCL was used as the template and amplified with pCas9-NG-F1 / pCas9-NG-R1, pCas9-NG-F2 / pCas9-NG-R2, and pCas9-NG-F3 / pCas9-NG-R3, respectively, to generate SpCas9-NG, the mutant of SpCas9 (Appendix Fig S1). These amplified fragments were then cyclized with the Gibson assembly method to construct pCas9-NG. To introduce a number of different gRNAs into a single cell, a high copy plasmid pSCM was used to load a number of different gRNA cassettes from the gRNA library (Appendix Fig S2). The specific process is as follows. pSCM was first used as the template and amplified with pSCM-F and pSCM-R to generate a kind of linearized plasmid skeleton. Second, pSCM was used as the template and amplified with the conventional forward primer gRNA-L-F and a reverse primer pool, gRNA-L-R, with 20 consecutive degenerate bases, to generate a random gRNA library containing about 4^20^ gRNA cassettes, the two flanks of which are over 50 bp homologous fragments to the two ends of the linearized plasmid skeleton, respectively. Third, both the linearized plasmid skeleton and the gRNA library were mixed together and delivered into the yeast cells by electroporation. Due to the adequate hybridity of gRNA cassettes in the gRNA library, a number of different gRNA variants can be randomly introduced into a single cell. With the aid of the powerful HR activity of *S. cerevisiae*, the linearized plasmid skeleton can be randomly cyclized with different gRNA cassettes in the yeast cells via the HR way. Due to the high copy number of pSCM, a number of different gRNA cassettes can be loaded into the cyclized pSCM plasmids. Only the cyclized plasmid skeleton with a specific gRNA cassette can be stable and generate gRNAs in the cells. Also because of the hybridity of gRNA cassettes, each cyclized plasmid may take a different RNA cassette with it during the cyclization process of pSCM and before their duplication in cell.

### Screening of high β-carotene producing strains

Standard preparations of competent cells and electroporation were used for the yeast transformation in this study (Kawai *et al*, 2010). First, appropriate amounts of gRNA library cassettes and plasmids skeletons were delivered into C0 competent cells and plated onto SD-Leu-Ura agar medium. Transformants were incubated at 30°C for 48 h. Colonies with darker orange colors were then inoculated into 5 mL of YPD medium and incubated at 30°C overnight, followed by inoculation into 30 mL of YDP and incubated on a shaker at 30°C for 72 h. The cells from each strain were collected to analyze β-carotene production and the evolved strain with highest β-carotene production was used for the next cycle of iterative evolution. After three cycles of iterative evolution, visual distinguishing of the darker orange colonies became difficult. Then, 96 deep well plates were used to culture and detect the further evolved strains, in which 96 single colonies were incubated in a 96 deep well plate with 0.5 mL of YPD medium at 30°C for 24 h to pre-select evolved strains with higher β-carotene production. Ten subsequently screened evolved strains were incubated in 30 mL of YPD on a shaker at 30°C for 72 h to re-select evolved strains that exhibited the highest β-carotene production. Overall, seven cycles of iterative evolution were completed over two months.

### β-carotene analysis

Standard β-carotene and lycopene compounds were dissolved in hexane and used to prepare standard curves. To quantify β-carotene production, cells were collected from 0.5 mL cultures and re-suspended in equal volumes of hexane. The re-suspended cells and 1.5 g of zirconia beads (0.5 mm diameter) were mixed in a 2 mL microcentrifuge tube and vibrated for 20 min at 60 Hz in a freeze grinder (Shanghai Jingxin, China) to disrupt cells and extract β-carotene. β-carotene measurements were performed during pre-screening by measuring absorption at 449 nm with a spectrophotometer (Thermo Labsystems Inc., PA, USA). Standard curves were generated to quantify carotenoids in the extracts. To re-screen the extracts, hexane containing the extracted carotenoids was filtered for HPLC analysis. A HPLC system equipped with an Agilent ZORBAX SB-Aq column (5 μm, 4.6 mm ×250 mm) and a UV/VIS detector was used to quantify β-carotene and lycopene levels at a wavelength of 474 nm at 30°C. The mobile phase used for gradient elution consisted of 90% aqueous acetonitrile (eluent A, HPLC grade) and methyl alcohol-isopropyl alcohol (3:2, v/v) (eluent B, HPLC grade). Gradient elution was performed at a constant flow rate of 1.0 mL/min with 100–10% eluent A and 0–90% eluent B for 0–15 min; 10% eluent A and 90% eluent B for 15–30 min; and 10–100% eluent A and 90–0% eluent B for 30–35 min (Zhou *et al*, 2015).

#### 2.6. Whole-genome sequencing and analysis

The genomes of the original strain C0 and the evolved strains C3-06, C5-63, and C7-143 were sequenced using PacBio single molecule real-time sequencing at Origingene (Shanghai, China). Extracted genomic DNA were prepared according to the manufacturer’s instructions for the PacBioz Template Prep Kit (Pacific Biosciences 10 kb template preparation protocol). The sequence data was filtered and the adapter sequences and low-quality data were removed, resulting in clean data that were used for subsequent analysis. The C0 genome was designated as the reference and genome variation information for samples was obtained by aligning sample reads against the reference. Finally, reads were mapped to the reference genome using the BWA software aligner and read coverage of the reference sequence along with alignment results were obtained using the SAMTOOLS software package. Structural variation (SV) referring to insertions (INS), deletions (DEL), inversions (INV), intra-chromosomal translocations (ITX), and inter-chromosomal translocations (CTX) between the reference and evolved genomes was identified by the Break-Dancer software package.

### gRNA randomness and number analysis

Genome editing was occurred within hours via CRISPR system (Liu *et al*, 2020b), however, it hard to immediately analyse the number of gRNAs in a single transformant after the electroporation. Considering that the types of gRNA would not change during the grow phase from a transformant to a clone in the plate, therefore, the number of gRNAs in a single clone grown from a transformant was analysed to represent the number of gRNAs in a single transformant. The clones with cyclized gRNA plasmids were selected with SD-Leu-Ura plates. To reduce error rates, ten clones were mixed together in equal levels for PCR amplification of the gRNA cassettes. The amplification of the random gRNA cassettes was conducted using 2 × PrimeSTAR^®^ Max DNA Polymerase (Takara, Beijing, China), 1 μmol of each of the forward and reverse primers (Appendix Table S3), and template in a 50 μL final reaction volume. PCR was performed as follows: 98°C / 3 min, 30 cycles of (98°C / 10 s, 60°C / 08 s, 72°C / 30 s) and a final extension of 72°C / 5 min. Subsequently, the PCR products were sequenced by Next-Generation sequencing with Illumina MiSeq at Personalbio (Shanghai, China), and three independent samples were sequenced to evaluate the average number of gRNAs in a single transformant.

### Transcriptional analysis

The original strain BY47419_NG_ as well as the C0 strain and the evolved strains C3-06, C5-63, and C7-143 were grown in YPD medium to exponential phase (OD_600_∼3) for transcriptomic analyses. A 50 mL aliquot of each yeast culture was then centrifuged at 8,000 × g for 2 min at room temperature. After removal of the supernatant, the cell pellets were frozen in liquid nitrogen, and total RNA was extracted using the QIAGEN RNeasy Mini kit. RNA sequencing via Next-Generation Sequencing was carried out with Illumina HiSeq at Personalbio (Shanghai, China). The KOBAS software package was used to investigate the statistical enrichment of differentially expressed genes among KEGG Pathways. Triplicate samples were used for transcriptional analysis.

### Statistical analysis

Unless otherwise stated, P-values for comparisons between conditions were estimated using an unpaired two-tailed *t-*test. Statistical details of the experiments can be found in the figure legends.

### Data access

The whole genome sequencing data generated in this study have been submitted to the NCBI BioProject database (https://www.ncbi.nlm.nih.gov/bioproject/) under accession number PRJNA682453.

The raw reads of the NGS data for the number analysis of gRNA-L in the evolutionary strains generated in this study have been submitted to the NCBI BioProject database (https://www.ncbi.nlm.nih.gov/bioproject/) under accession number PRJNA680435.

The transcriptomic data generated in this study have been submitted to the NCBI Gene Expression Omnibus (GEO; https://www.ncbi.nlm.nih.gov/geo/) under accession number GSE162126.

## Acknowledgments

This work was financially supported by the Natural Science Foundation of Shanghai (No.20ZR1415100) and the Chinese Postdoctoral Science Foundation (No. 2020M671021).

**Expanded View** for this article is available online.

## Author contributions

MZ, FW, and DW designed the research. MZ performed CRAM construction experiments, strains evolution experiments. MG performed growth curves, β-carotene analysis, laser scanning confocal microscopy, and cloning. LX and YL assisted with cloning and preliminary experiments. MZ, XT, BG, and ML analyzed the results. DW supervised the research. MZ and FW wrote the manuscript.

## Conflict of interest

The authors declare no competing interests.

